# The phase of tACS-entrained pre-SMA beta oscillations modulates motor inhibition

**DOI:** 10.1101/2023.08.25.554760

**Authors:** Zhou Fang, Alexander T Sack, Inge Leunissen

## Abstract

Inhibitory control has been linked to beta oscillations in the fronto-basal ganglia network. Here we aim to investigate the functional role of the phase of this oscillatory beta rhythm for successful motor inhibition. We applied 20 Hz transcranial alternating current stimulation (tACS) to the pre-supplementary motor area (pre-SMA) while presenting stop signals at 4 (Experiment 1) and 8 (Experiment 2) equidistant phases of the tACS entrained beta oscillations. Participants showed better inhibitory performance when stop signals were presented at the trough of the beta oscillation whereas their inhibitory control performance decreased with stop signals being presented at the oscillatory beta peak. These results are consistent with the communication through coherence theory, in which postsynaptic effects are thought to be greater when an input arrives at an optimal phase within the oscillatory cycle of the target neuronal population. The current study provides mechanistic insights into the neural communication principles underlying successful motor inhibition and may have implications for phase-specific interventions aimed at treating inhibitory control disorders such as PD or OCD.

## Introduction

Motor inhibition is a crucial cognitive function that allows us to rapidly adapt to our ever-changing environment by cancelling or suppressing planned movements. Previous studies have found that this essential inhibitory control is associated with the activation of a fronto-basal ganglia network, including the pre-supplementary motor area (preSMA), the right inferior frontal cortex (rIFC), and the subthalamic nucleus (STN) (Jahanshahi et al., 2015). These key nodes of the fronto-basal ganglia inhibition network are presumed to interact with each other via the so-called “hyperdirect pathway” (Aron, 2007; Aron et al., 2014; Jahanshahi et al., 2015).

There is concrete evidence suggesting that this inter-regional information exchange is directly related to beta-band oscillatory mechanisms within the fronto-basal ganglia circuit. Intracranial recordings from these regions revealed increased oscillatory activity in the beta frequency band during successful inhibition (Alegre et al., 2013; Kühn et al., 2005; Swann et al., 2009; Swann et al., 2012; Wagner et al., 2018; Wessel and Aron, 2013; Wessel et al., 2016; Castiglione et al., 2019), and this increase in beta activity relates to inhibitory performance (Schaum et al., 2021; Leunissen et al., 2022; Ding et al., 2023). Moreover, the hyperdirect pathway is overactive in Parkinson’s disease (PD) (Jahanshahi et al., 2015), resulting in excessive beta oscillations in the STN and cortical motor areas (Brown, 2007), which are associated with bradykinesia, rigidity, and freezing symptoms (Kühn et al., 2008; Little et al., 2012; Chen et al., 2010; Toledo et al., 2014). Finally, entraining beta oscillations by means of transcranial alternating current stimulation (tACS) promotes inhibition of unintended movements (Joundi et al. (2012); Leunissen et al., 2022; Tan et al., 2023), suggesting that the power (‘amplitude”) of beta oscillations is causally related to successful motor inhibition. However, although the functional implications of beta oscillations for motor inhibition have been demonstrated in previous research, not much is known yet about the exact temporal dynamics of neural communication within the fronto-basal ganglia circuit.

Long-range neural communication is thought to come about through groups of neurons engaging in rhythmic synchronization, creating short temporal windows of low and high excitability of the respective regions (Fries, 2005). Effective neural communication between such regions requires a stable phase relation between sending and receiving populations, known as coherence. The disruption of beta phase coherency between the STN and motor/premotor regions by deep brain stimulation suggests that it is in fact the phase of oscillatory beta that may play a crucial role in successful inhibitory control (Oswal et al., 2016; Holt et al., 2019; Salimpour et al., 2022). In line with this notion, local broadband neuronal activity in the motor cortex seems to be entrained on the phase of the beta rhythm (Miller et al., 2012) and motor cortex excitability is beta-phase dependent (van Elswijk et al., 2010; Keil et al., 2013; Khademi et al., 2018; Torrecillos et al., 2020; Wischnewski et al., 2022).

Conventional methods for investigating oscillatory phase, such as magneto-/electroencephalography, are limited by their post-hoc nature and by providing only correlational evidence. This is particularly problematic when investigating stimuli that require a psychophysical staircase, such as the stop signal delay in a stop signal task which is typically staircased to maintain a 50% stopping accuracy to allow for an estimation of stopping speed (Verbruggen et al., 2019). In contrast, tACS can phase align cortical neural activity with the externally applied current (Thut et al., 2011; Herrmann et al., 2016), transforming the phase of oscillatory brain activity into an experimentally controlled independent variable (ten Oever et al., 2016). In the current experiment we applied beta tACS over the preSMA to experimentally control the phase of oscillatory beta activity while presenting behaviorally relevant stop signals precisely time-locked to 4 (experiment 1) or 8 (experiment 2) pre-determined equidistance phases of the tACS waveform. Stop signal delay staircases were run in parallel for the different phase conditions in order to investigate the causal role of the beta phase for successful motor inhibition. Assuming that the fronto-basal ganglia network makes use of the beta rhythm to convey the need for inhibition between the pre-SMA / frontal regions and the subcortical downstream nodes of the fronto-basal ganglia network, we expected that stop signals presented at specific and distinct beta phases would lead to improved inhibitory control whereas other beta phases may cause decreased motor inhibition performance.

## Results

### Experiment 1

All participants (N = 32) of experiment 1 tolerated the stimulation well as indicated by the VAS fatigue and discomfort rating score (Table S1). Although there is a significant change between pre-and-post fatigue scores, the overall fatigue and discomfort levels are within the mild range. Three participants were excluded from further analyses. Data for one participant was lost due to a technical issue; another participant had an average stop rate >70%; and the last participant violated the context independence assumption, resulting in a total of 29 participants.

#### Behavioral results

A repeated measures ANOVA was used to compare the effect of phase on SSRT, RTsf, and P(inhibition), and a Friedman test was performed on force peak and force peak rate. The results show that all behavioral outcomes differ significantly between phases (Table 1). Post-hoc tests revealed a significant difference in both SSRT and P(inhibition) between 90°and 270°-360°, and 180°and 270°-360°phase conditions (p<0.05); in RTsf between 270°and 90°-180°-360°, and 180°and 270°-360°(p<0.05). Wilcoxon Signed Rank test results show a significant difference in both the peak force and peak force rate of stop trials, between 90°and 270°-360°, and 180°and 270°-360°phases (p<0.05) (Table S5, S6). The post-hoc analysis yielded compelling results indicating that subjects exhibited significantly higher accuracy and faster response times when stop signals were presented at 270°-360°compared to 90°-180°. Furthermore, the reaction times on fail stop trials were the shortest at 270°, reflecting that only the fastest go processes could overcome the inhibition at this phase. The Wilcoxon Signed Rank test further revealed that participants exerted less force and lower force rate on stop trials at 270°-360°compared to 90°-180°, indicating a significant increase in inhibitory control. In conclusion, the findings highlight a greater degree of inhibition at the trough phase, 270°-360°.

**Table 1.**
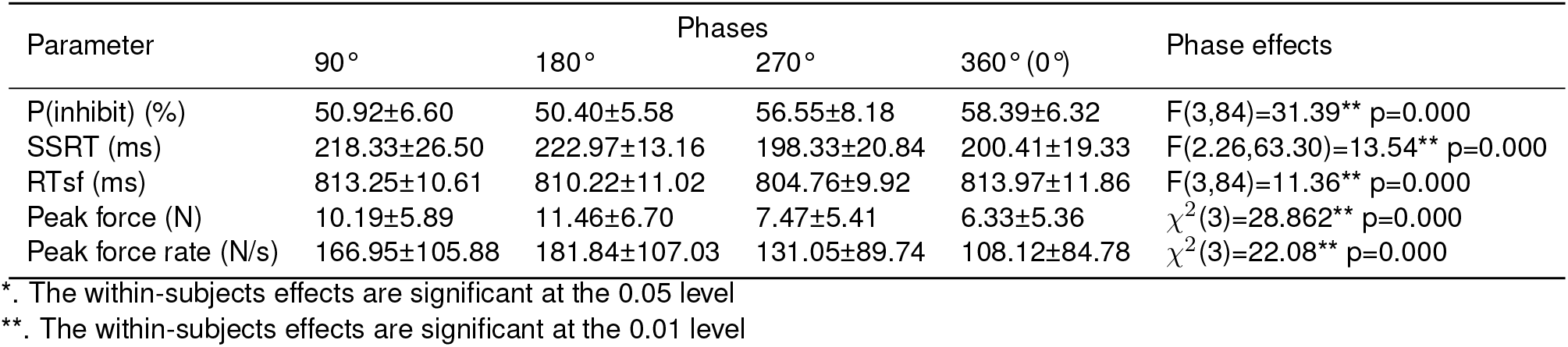
Group results. The phase effects with p-value were calculated from repeated measures ANOVA.

Figure 1 shows the group results from Exp.1 across 4 tACS-phase conditions, superimposed with the best-fitting one-cycle sinusoids. In line with the results of the ANOVA, the curve-fitting results also show a significant phase modulation of behaviour that matches a one-cycle sinusoid. However, the interpretation of these results warrants some caution as a sinusoidal curve fitting with only four data points can lead to inaccurate outcomes. This led to Exp.2 where we increased the phase conditions to 8.

**Figure 1.**
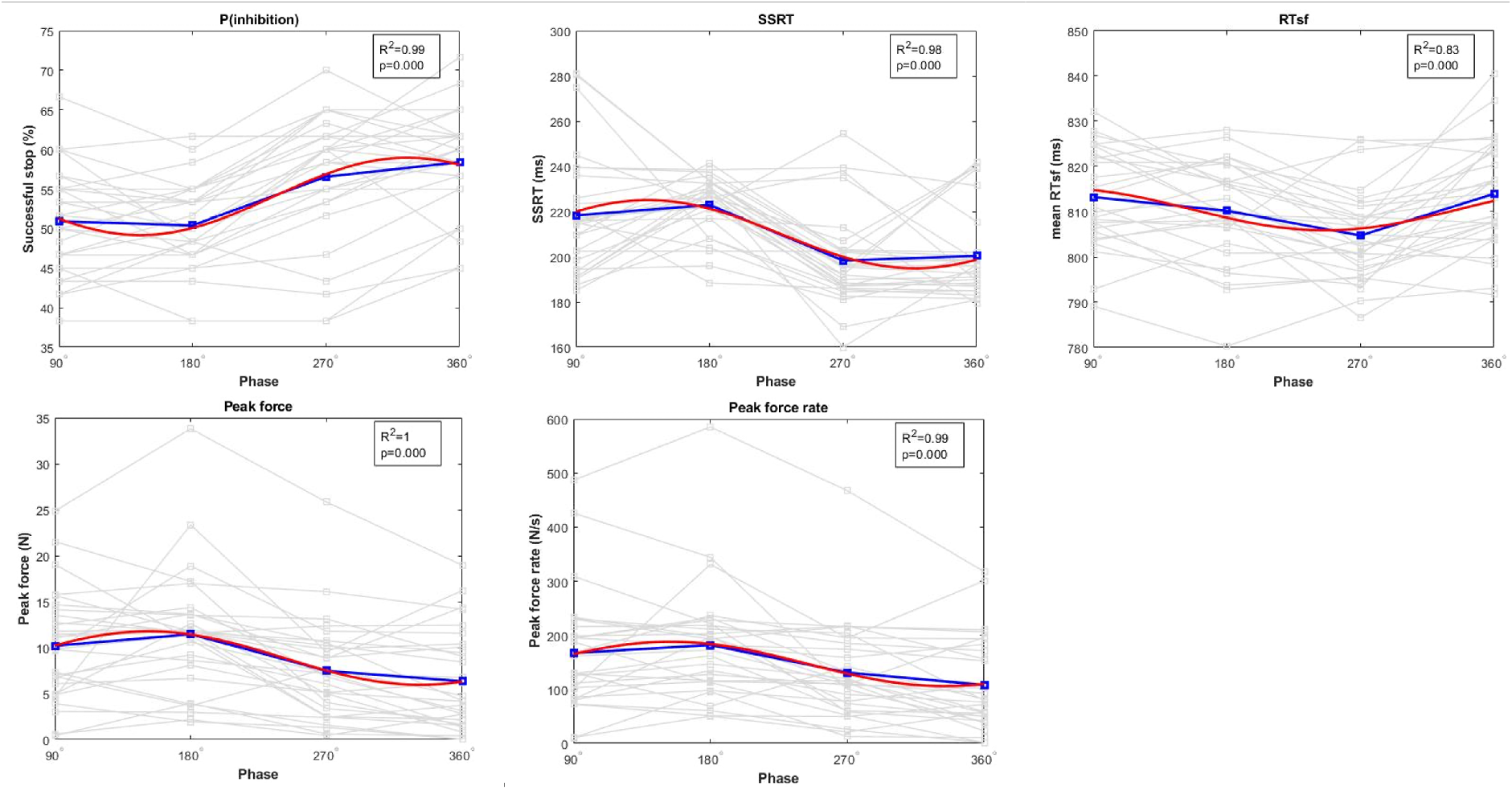
Group behavioral results of Exp.1. **P(inhibition)**: the percent of successful stops; **SSRT**: stop signal reaction time; **RTsf**: reaction time on failed stops; **Peak force**: the maximum force; **Peak force rate**: the maximum rate of change in the force signal. The individual behavioral outcomes are shown as grey lines. The group average value is in blue, and the best-fitting sinusoid is in red. In the legend boxes, the R^2^, which indicates the goodness of fit, along with the p-value based on the permutation test on the relevance value are provided.

Although similar patterns of phase dependency were observed for all participants, with a sine wave shaped relation, the best inhibitory performance does not necessarily occur at the same beta phase in each participant. In the event that the optimal phase varies between participants, averaging data in each phase condition might conceal an existing phase dependency on the group level. To this end, the peaks of individual subject data were re-aligned to the 90°phase before averaging across participants. Such realignment creates an artificially positive modulation at the optimal phase, therefore the data were tested against shuffled data that underwent the same peak realignment procedure. The results of this phase-aligned analysis can be found in the supplement (Figure S1).

### Experiment 2

In Exp.2, the VAS fatigue and discomfort scores indicated that all the participants (N=16) tolerated the stimulation well in both sessions (Table S2). Repeated measures ANOVA shows no difference between the two sessions. The participant that had a successful stop percentage >70% and the one that was lost due to the technical issue were excluded from further analyses. As a result, 14 participants remained.

#### Behavioral results over phases

We ran a repeated measures ANOVA to determine if there was a statistically significant difference in SSRT, RTsf and percent of successful stops between each of the eight phases. For the non-normally distributed measures, peak force and peak force rate, we used a Friedman test. The statistic results are shown in Table 2. All measures show significant effects from phase conditions. Post hoc results show a significant change in P(inhibition) between 90° and 360°, 180° and 270° -360°; in SSRT between 135° and 270°; in RTsf between 270°and 90°-135°(Table S7); Wilcoxon Signed Rank test results indicate that in peak force, there is a significant difference between 270°and 90°-135°-180°-360°, 315°and 90°-135°-225°, 360°and 90°-135°-180°-270°-225°; in peak force rate between270°and 180°-225°-360°, 315°and 180°-225°, 360°and 90°-135°-180°-270°(Table S8). Aligned with the findings of Exp 1, the post hoc analysis and Wilcoxon Signed Rank test consistently show that 270°-360°elicits the highest accuracy in successful stops and the shortest reaction times on stop trials. Notably, at 270°, the reaction time on failed stop trials was significantly reduced. Additionally, at 270° -360°, both force and force rate were significantly lower. These converging findings demonstrate that participants displayed better inhibition when stop signals were at the trough phases of beta oscillations.

**Table 2.** Behavioral variances across 8 phase conditions, and the effects of phase modulation effects resulting from the repeated measures ANOVA.

| Parameter | 45° | 90° | 135° | 180° | 225° | 270° | 315° | 360° | Phase effects |
| --- | --- | --- | --- | --- | --- | --- | --- | --- | --- |
| P(inhibition) (%) | 53.40±6.60 | 50.83±7.18 | 51.25±8.79 | 50.48±6.49 | 50.24±13.30 | 55.60±8.61 | 53.21±10.30 | 59.29±7.70 | $F(3.42,44.46)=4.03^{**}$<br>$P=0.001$ |
| SSRT (ms) | 211.46±24.73 | 219.29±29.84 | 233.89±28.97 | 220.89±15.04 | 225.14±41.95 | 197.00±20.79 | 207.61±34.90 | 200.36±19.56 | $F(7,91)=3.39^{**}$<br>$P=0.003$ |
| RTsf (ms) | 812.47±13.41 | 814.92±12.14 | 816.87±11.65 | 810.92±13.14 | 810.07±14.91 | 804.53±10.13 | 812.76±13.60 | 814.12±14.15 | $F(7,91)=2.94^{**}$<br>$P=0.008$ |
| Peak force (N) | 9.77±6.23 | 9.93±5.90 | 11.12±6.19 | 11.25±6.35 | 10.85±7.33 | 7.87±5.72 | 8.22±5.15 | 5.94±5.14 | $\chi^2(7)=26.31^{**}$<br>$p=0.000$ |
| Peak force rate (N/s) | 148.19±76.99 | 161.11±73.47 | 163.45±77.43 | 184.64±79.59 | 168.44±93.25 | 139.18±74.82 | 132.36±69.14 | 103.62±66.45 | $\chi^2(7)=24.79^{**}$<br>$p=0.001$ |

\*. The within-subjects effects are significant at the 0.05 level
\*\*. The within-subjects effects are significant at the 0.01 level

Then we looked at the causal role of the tACS phase on inhibition. Figure 2 depicts the group results of behavioral outcomes over eight phase conditions (blue plot), with best-fitting sinusoids in red. The fitted sinusoids reveal significant effects on P(inhibition) (R^2^=0.61), SSRT (R^2^=0.75), RTsf (R^2^=0.59), peak force (R^2^=0.42) and peak force rate (R^2^=0.64), p<0.05, showing that these measures are highly dependent on the phase where inhibition occurs. These results indicate that inhibitory performance over tACS-phases matches a sinusoidal oscillatory pattern, and inhibition is at its greatest when the stimuli are presented at the trough of the tACS oscillation, which is in line with the results of Exp. 1.

**Figure 2.**
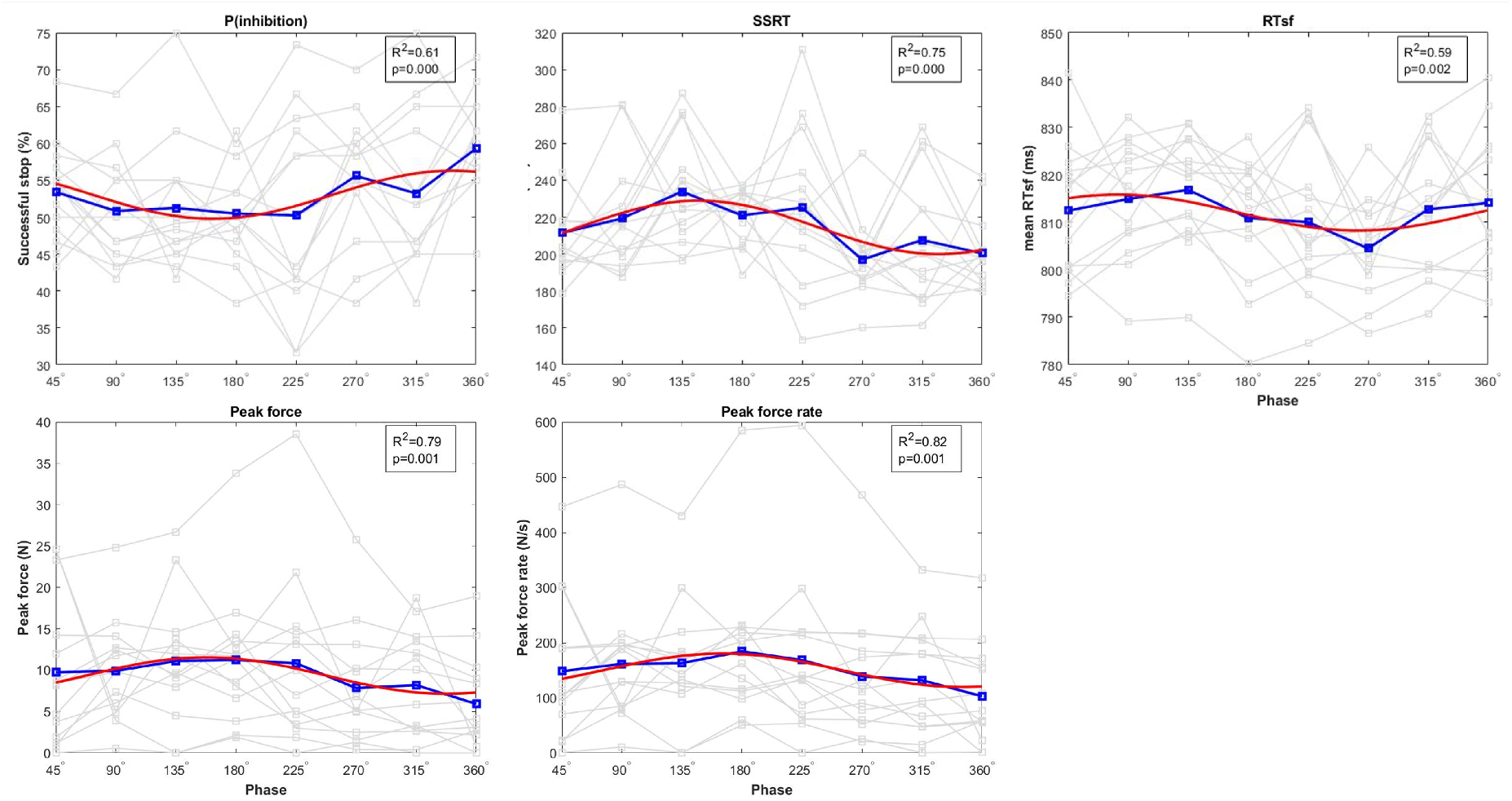
Group behavioral results of Exp.2. **P(inhibition)**: the percent of successful stops; **SSRT**: stop signal reaction time; **RTsf**: reaction time on failed stops; **Peak force**: the maximum force; **Peak force rate**: the maximum rate of change in the force signal. The individual behavioral outcomes are shown as grey lines. The group average value is in blue, and the best-fitting sinusoid is in red. In the legend boxes, the R^2^, which indicates the goodness of fit, along with the p-value is provided.

#### EEG results

All results are based on the signal recorded at Fz as it is located over the preSMA and neighbours Fcz which is the central tACS electrode.

### Pre- and post-resting EEG

We conducted a paired t-test to compare the absolute power of the pre- and post-beta (15-30 Hz) band during the resting state, averaged over participants in each session (N=28) (Figure 3 A). The results revealed a significant increase after stimulation (0.051±0.073) compared to before (0.015±0.008), t(27)=2.573, p=0.016. The results indicated that the overall rest-state beta power significantly increased after the experiment session.

**Figure 3.**
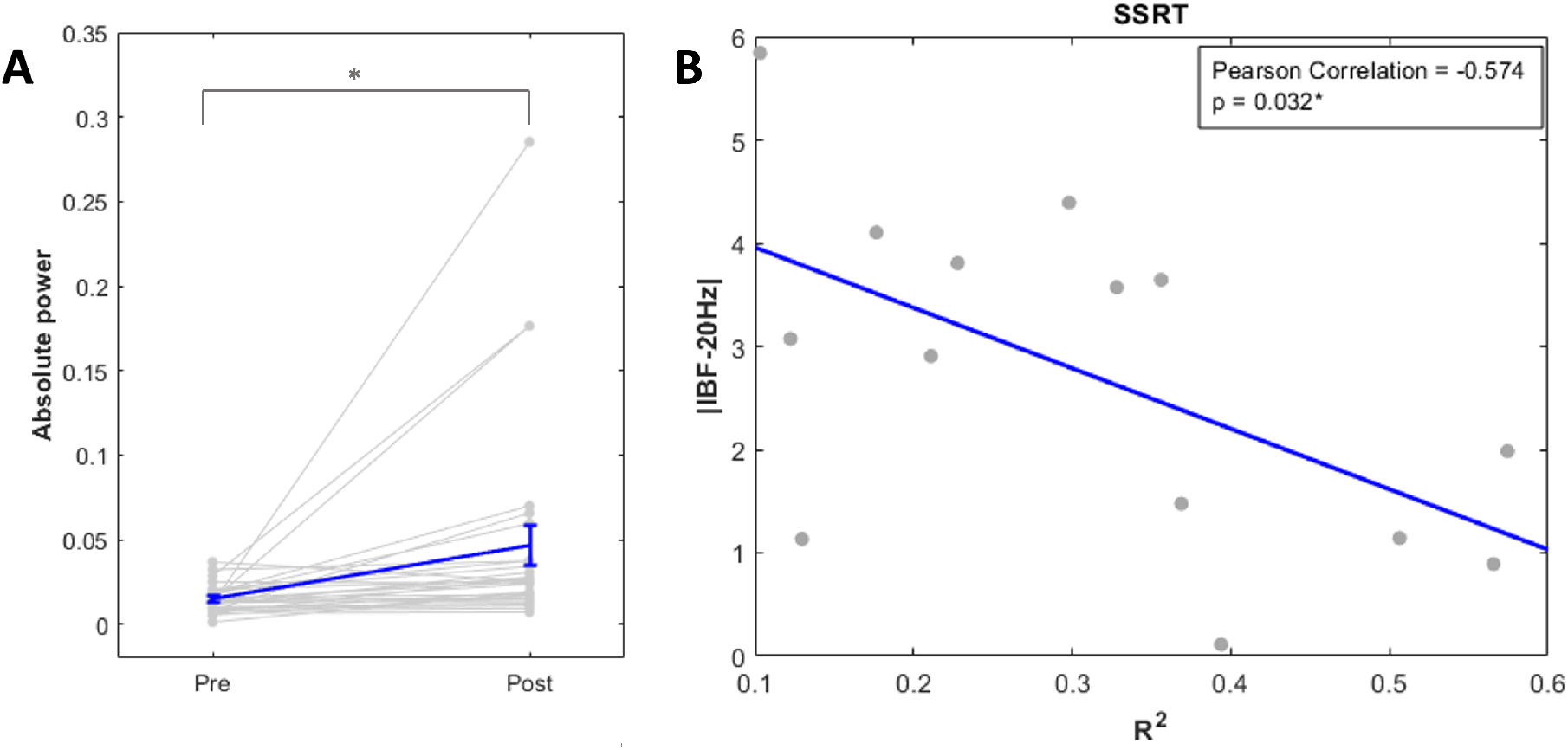
EEG results. A) Pre-post beta absolute power with 95% confidence interval error bars. B) Correlation between R^2^ of measures and distance between IBF and 20 Hz.

### Individual beta peak frequency

When aligning stimuli to the tACS waveform, it is crucial that the tACS entrains the endogenous oscillations for one to be able to observe a phase modulation in behaviour. It is likely that the strength of entrainment varies between individuals. Based on previous studies, demonstrating that 20 Hz is the central beta frequency in the motor system (Chakarov et al., 2009; Muthukumaraswamy, 2010; Fischer et al., 2017), we set the tACS stimulation frequency as 20 Hz over the pre-SMA area to modulate inhibition behaviour. According to the Arnold tongue principle, entrainment happens when the external stimulation frequency is close to the intrinsic oscillation frequency and/or the stimulation intensity is sufficiently high (Ali et al., 2013). To investigate whether the phase modulation effects were stronger when the endogenous oscillations were closest to 20Hz, we correlated the individual beta frequency based on the pre-resting state EEG of each participant in Exp 2 with the R^2^ of their inhibition performance variables (Figure 3 B). The Rsq of SSRT, peak force and peak force rate showed a significant correlation with the distance between IBF and stimulation frequency (20 Hz) (SSRT: r=-0.574, p=0.032; Peak force: r=-0.696, p=0.006; Peak force rate: r=-0.561, p=0.037), indicating that the closer the frequency of the endogenous brain oscillation to the external stimulation, the stronger the phase modulation of behavior.

## Discussion

The aim of the current study was to investigate the functional role of the oscillatory beta phase within the fronto-basal ganglia network for successful motor inhibition. We presented inhibition stimuli at various pre-determined and experimentally controlled phases of tACS entrained beta oscillations. This methodology was achieved with the setup developed by ten Oever et al. and previously used in phase-dependent studies by de Graaf et al. (2020) and Schilberg et al. (2018). Using this setup, we experimentally aligned the stop signal onset with the 4 (Exp 1) and 8 (Exp 2) pre-determined phases of the entraining tACS oscillation. Successful inhibitory control at each of these phases was assessed by participants’ performance in the stop signal task, where their stop-signal reaction time (SSRT), successful stop rate (P(inhibition)), the reaction time of stop fail (RTsf), peak force and peak force rate in stop trials were tracked. We found that inhibition performance was systematically affected by the beta phase at which stop signals were presented. Moreover, aligned with our a priori hypotheses, group results showed that inhibition outcomes over these selected phases fit a sinusoidal pattern. Specifically, better inhibitory control was observed at the trough of the beta phase whereas inhibition control for stop stimuli presented at the peak was significantly decreased.

### Effect of beta oscillatory phase on inhibitory motor control

Previous studies have consistently shown that successful motor inhibition is associated with increased beta power and beta bursting in both frontal and subcortical nodes of the motor inhibition network (Swann et al., 2012; Ray et al., 2012; Wessel and Aron, 2013; Alegre et al., 2013; Fonken et al., 2016; Schaum et al., 2021; Diesburg and Wessel, 2021), suggesting that the fronto-basal ganglia circuit fine-tunes inhibitory motor control by adequately regulating beta oscillatory network activity. More conclusive evidence for a causal link between beta oscillatory activity and motor inhibition stems from transcranial electric stimulation studies demonstrating that increasing the amplitude of beta oscillations in the motor system results in stronger motor inhibition (Joundi et al., 2012; Leunissen et al., 2022; Tan et al., 2023).

However, to fully understand the dynamics within this circuit during motor inhibition we must consider the role of oscillatory beta phase. Phase coherence between brain areas is believed to facilitate effective communication by creating windows in which information can be transmitted most efficiently (Fries, 2005; Fries, 2015). Delivering stimuli during an optimal phase should therefore enhance behavioral performance. Such a relationship between the phase-coupled presentation of stimuli and performance has been shown in other cognitive domains such as sensory perception (Bush et al., 2010; Kasten and Herrmann, 2020), attention (VanRullen, 2018) and memory (Ten Oever et al., 2020). With respect to motor inhibition, stronger coherence in the beta band has been associated with faster inhibitory control (Swann et al., 2012; Ding et al., 2023). However, until now, it was unclear whether inhibitory control performance functionally depends on the phase at which the stop signal is presented.

Our study provides causal evidence for a modulatory role of the beta oscillatory phase on motor inhibition that follows a one-cycle sinusoidal pattern. Specifically, we discovered that inhibitory control performance was improved for stop signals presented at the trough phase of the experimentally controlled oscillatory beta wave, while inhibition performance was decreased at the peak of the beta phase. This relationship was systematic within participants and surprisingly consistent across participants. This consistent relation between experimentally controlled beta-phase and inhibitory control performance supports the notion that input arriving at the excitatory phase of the local neuronal ensemble increases the efficiency of information transfer when task-induced coherence connects the cortical neural population (preSMA) to the functional brain network (Pérez-Cervera et al., 2020; Schneider et al., 2021). The degree of consistency in this systematic phase-dependency indicates that the optimal phase relation is stabilized and brought about by the enduring anatomical structure of the network pathway (Deco and Kringelbach, 2016).

The beta-phase-dependent inhibitory control in this study corroborates the findings that motor cortex excitability varies with the beta oscillatory phase. Miller et al. (2012) demonstrated that the beta rhythm acts as a suppressive mechanism to actively gate local motor cortex activity, with the highest activity on the falling flank of the beta cycle. This also aligns with the finding of Hussain et al. (2022) that motor commands are released between 120°-140°along the beta cycle, and that TMS stimulation on the falling flank of the beta wave results in the highest motor evoked potentials (MEP), whereas stimulation on the trough or rising flank results in smaller MEPs (Torrecillos et al., 2020; Wischnewski et al., 2022, but note that different phases have been reported as well: Schilberg et al., 2018; van Elswijk et al., 2010; Khademi et al., 2018). Interestingly, Cagnan et al. (2019) found that action potentials in the STN are typically phase locked to the 270°phase of frontal beta oscillations. In addition, GABAa-dependent intracortical inhibition in M1 is largest at the 270°phase (Guerra et al., 2016). Together, these findings suggest that (dis)inhibitory signals can be transmitted from the frontal cortex to M1 via the basal ganglia – thalamo circuits at beta frequency with an optimal window for motor inhibition and initiation at opposing phases of this oscillatory beta cycle.

Our results highlight the role of the beta phase in neuronal hypo-or hyper-synchronization where (de)synchronization within brain networks helps to weaken or facilitate communication in a phase-specific manner. Interfering with a specific phase could therefore disrupt dysfunctional communication resulting from abnormal synchronization in neurological and psychiatric disorders. Excessive synchronization of beta oscillations attributed to dopamine deficiency in PD has for example been observed across cortical areas and the BG (Williams et al., 2002; Litvak et al., 2010; Hirschmann et al., 2011; Koren et al., 2022), and disrupting beta synchronization in cortical (Doyle Gaynor et al., 2008) and BG nodes, including striatum (Costa et al., 2006; McCarthy et al., 2011), pallidum (Kühn et al., 2008), subthalamic nucleus (de Solages et al., 2010; Darcy et al., 2022) and thalamus (Van Der Werf et al., 2006), results in improvements of bradykinesia, rigidity, and freezing. Precisely timing transcranial magnetic stimulation (TMS) to the opposing phase of the optimal window of ongoing beta oscillations in cortical regions may likewise present an effective non-invasive intervention for personalized modulation (Cassidy et al., 2002; Sharott et al., 2018), similar as has been demonstrated for phase-specific STN stimulation (Holt et al., 2019).

### Phase modulation in relation to tACS entrainment

By employing tACS and pre-programming the presentation of the stop signals at specific phases of the tACS wave, we were able to conduct psychophysical staircases over each of the experimentally controlled phase conditions. However, one crucial factor in studying phase modulation with tACS is the entrainment of cortical areas to external stimulation. The pre-post EEG comparison shows a significant increase in beta power suggesting a potential aftereffect of the stimulation, which could indicate successful entrainment. However, due to the absence of a sham condition, we cannot rule out the possibility that the observed increase in beta power might be attributed to task performance, rather than solely tACS entrainment. The Arnold tongue principle illustrates that external stimulation with a frequency close to the underlying internal frequency in the cortical area leads to the highest entrainment (Ali et al., 2013) (i.e., highest phase-locking between endogenous and exogenous oscillations with a consistent phase difference (Negahbani et al., 2019; Vogeti et al., 2022)), and this entrainment is dose-dependent (Johnson et al., 2020). In our study, we revealed a significant correlation between the distance of the individual peak frequency from the 20 Hz stimulation frequency and the degree of phase modulation of inhibitory control performance (r=0.574, p=0.032), indicating that better entrainment with tACS led to a more sinusoidal pattern of inhibitory control performance. This phenomenon is likely attributed to the strong alignment between endogenous beta oscillations and the tACS waveform. Consequently, stop signals, synchronized with the phase of the tACS waveform, reached the desired phase of the intrinsic brainwave with greater precision.

## Conclusion

The current study provides evidence for a functional modulatory role of the oscillatory beta phase within the fronto-basal-ganglia network for inhibitory motor control performance. We revealed that the trough of experimentally controlled beta oscillations in the preSMA provides an optimal window for exerting inhibition control over presented stop signals. In addition, participants with an individual beta peak frequency close to the 20Hz tACS stimulation frequency showed the strongest phase-dependent modulation of inhibitory motor control performance, further supporting the notion that the phase of beta oscillations is casually involved in successful motor inhibition. These findings provide insights into the mechanism of neural communication underlying motor inhibition and add to the existing knowledge of phase dependency in the motor system. The obtained results may guide the development of phase-locked non-invasive stimulation treatments for inhibitory control disorders such as PD and OCD.

## Methods and Materials

To investigate the relationship between the oscillatory phase and inhibitory performance, we conducted two related experiments (Exp 1 and Exp 2), with Exp 2 adding four more conditions on top of Exp 1. The following section describes the experimental procedures of Exp 1 and 2 as well as how they differ.

### Participants

Thirty-two healthy, right-handed (the mean laterality quotient score is 96.3, with a range of 70–100 (Oldfield, 1971)) 18-35 year-old (mean age 25.3; 13 males and 19 females) volunteers were recruited for this study. All 32 participants were involved in Exp 1 and 16 of them took part in both Exp 1 and Exp 2 (mean age 24.3; 8 males and 8 females). Standard screening verified that there were no contraindications to non-invasive brain stimulation (Bikson et al., 2009; Woods et al., 2016). All procedures were approved by the local ethical committee and written informed consent was obtained from all participants.

### General experiment procedure

In all experiments, participants received 20 Hz (beta) tACS stimulation during the performance of a stop-signal task. In Exp 1, the stop signals were presented at 4 equidistant phases of one oscillatory cycle of the tACS signal: 90°, 180°, 270°, and 360°(0°). The procedure of Exp 2 was split into two days more than 48 hours apart, with four phase conditions completed each day: day 1: 90°, 180°, 270°, and 360°(0°) and day 2: 45°, 135°, 225°, and 315°. The order of the two sets of conditions was counterbalanced across participants.

At the beginning of each experimental session, a 3-minute resting EEG was recorded while participants kept their eyes open and fixed on a black screen. Subsequently, participants practised the stop signal task by performing 20 go trials, followed by 20 trials in which go and stop trials were mixed and next performed four experimental blocks, each including 180 trials. Therefore, Exp 1 contained 720 trials per subject, and Exp 2 had a total of 1440 trials. Participants had a 3-5-minute break between each block. After the task, the post-EEG was recorded in the same way as the pre-EEG.

Participants were asked to report their self-perceived level of fatigue before and after the session, and their tACS discomfort level after the experiment on a Visual Analogue Scale (VAS). The VAS consisted of a 10 cm horizontal line with two opposing limits displayed at either end of the line (no discomfort-worst discomfort for the discomfort level and not tired-exhausted for the fatigue level).

### Stop-signal paradigm

Participants were asked to sit comfortably, 1 meter away from the screen (refresh rate 240 Hz) and perform an anticipated response stop-signal task (Slater-Hammel, 1960; Coxon et al., 2006; Leunissen et al., 2017) as shown in Figure 4 A. The stop-signal task was programmed in LabVIEW (National Instruments, Austin, TX). Each trial started with the display of an empty bar in the centre of the screen, with an indicator filling it over the course of 1000 ms at a constant speed from bottom to top. A target line was positioned at 800 ms from the onset. Participants were instructed to pinch a force transducer (OMEGA Engineering, Norwalk, CT, USA) with their right index finger and thumb to stop the indicator when it reached the target line. A force response threshold was initially set at 25% of their maximal voluntary force (MVF), which could be lowered in 5% increments until participants were able to comfortably reach the threshold during the whole session (mean is 23.3%; range is 10% 25%). MVF was determined as the highest force value measured during 3 maximal strength pinches of 5s. Participants were encouraged to stop the indicator at the target line as accurately as possible. The target line changed colour at the end of the trial as feedback on their performance. There were 4 colour options: green, yellow, orange, and red, indicating the distance between the indicator and the target line within 20, 40, 60, and >60 ms, respectively. 33% of the trials were stop-signal trials, in which the indicator automatically stopped before reaching the target line. In this case, participants had to refrain from pinching the sensor. Throughout stop-signal trials, the indicator was equally likely to stop at 110 ms, 150 ms, 190 ms, 230 ms, and 270 ms before the target line. The timing was slightly shifted (maximally ±25 ms) to match the closest desired phase of the tACS wave. After 1s the indicator was reset to empty. The inter-trial interval jittered between 2.7-3.7 ms. Participants completed four task runs per session, each comprising 120 go and 60 stop trials presented in a pseudorandomized order (720 trials per session in total). Each task run contained stop signals presented at one of the four desired phases at all possible stop-signal delays. For more information on the synchronization between tACS wave and stop-signal presentation see section: tACS phase locked stop-signal presentation.

**Figure 4.**
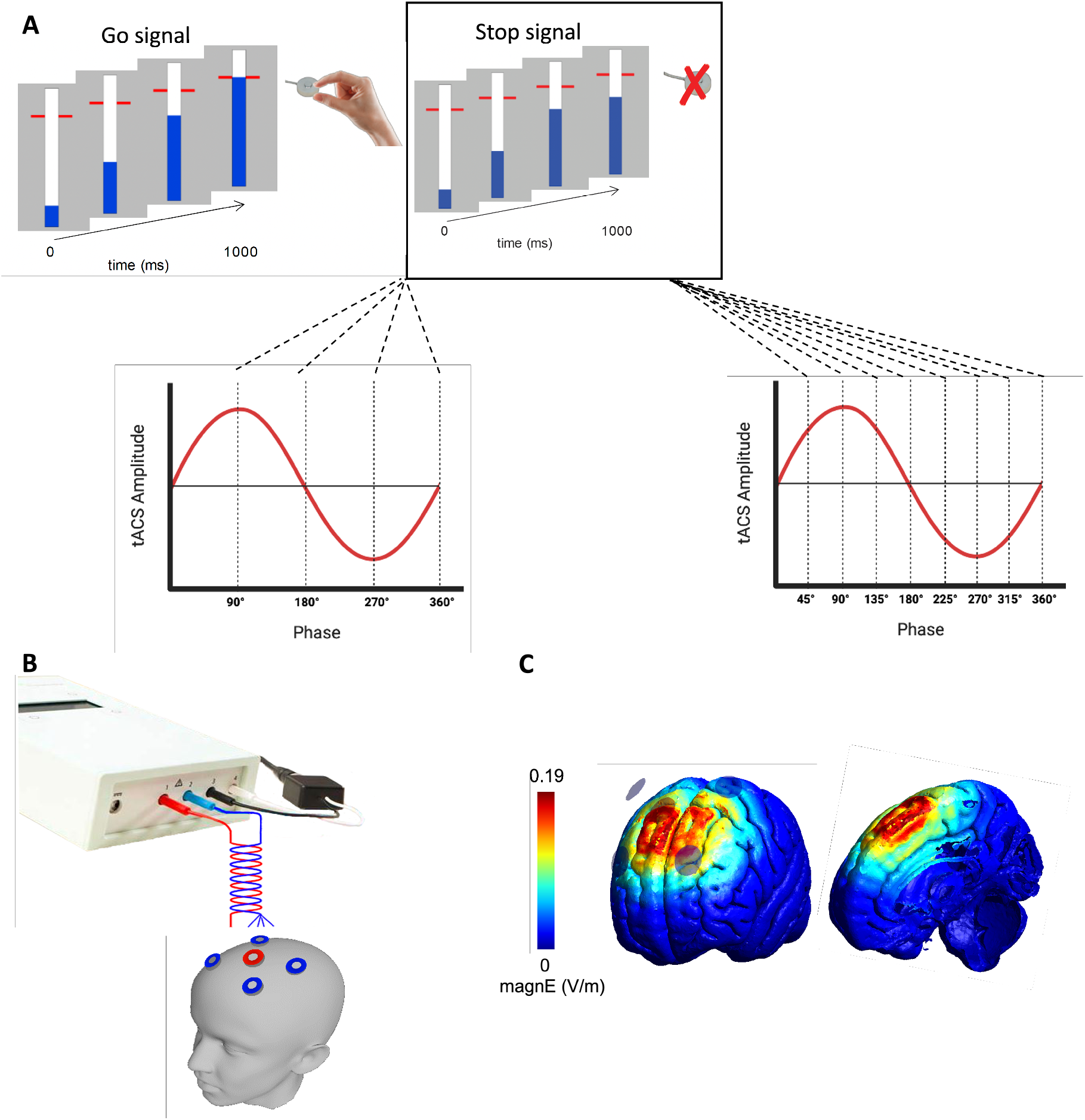
Experimental methods. A) Stop-signal paradigm. The blue indicator filled the empty bar at a constant speed. In go trials, it rose from the bottom to the top and participants were asked to stop it at the target line. In stop trials, the indicator stopped increasing before the target line. In this case, participants had to withhold their responses. Stop signals were aligned with the 4 (Exp 1) and 8 (Exp 2) desired phases of the tACS wave. B) tACS electrode montage. The centre electrode ring is placed over FCz and surrounded by four electrode rings at a centre-to-centre distance of 5 cm. C) Simulated electrical field distribution generated by the tACS setup in an example brain.

The force signal was captured through a digital-analog converter (DAC) from National Instruments (Austin, TX, USA) and saved trial-by-trial (from the indicator starting filling up to 1700 ms) using LabChart with a sampling rate of 1000 Hz. Stop trials were classified as failed stop trials if the force produced exceeded the response threshold. If the force remained below the threshold, the trial was classified as successfully inhibited.

### tACS procedures

20 Hz tACS was applied using a 4 × 1 HD-tACS setup (DC STIMULATOR PLUS, NeuroConn, GmbH, Ilmenau, Germany). Custom made gel-filled cup-electrodes made out of plastic cylinders (∅2cm at top, 2.5cm at bottom) were mounted in an EEG cap (EASYCAP GmbH, Germany) and filled with electrode gel (OneStep Cleargel) before an EEG Ag/AgCl ring electrode was fastened into the cup. The target electrode was placed over the preSMA (FCz) and four surrounding electrodes were positioned at F1, F2, C1, and C2 based on the international 10-20 coordinate system (Figure 4 B). The impedance of the tACS electrodes was kept below 10 kΩ (8.4±1.6 kΩ) and the peak-to-peak amplitude of the stimulation was set to 2 mA by default, which was reduced if the participant were too uncomfortable with the stimulation resulting in the mean intensity being 1.97±0.11 mA. The stimulation was ramped up and down for 10 s and lasted 9.6 min in total per block leading to a 38.4 min total stimulation received by the participants per session. The electrical field distribution generated by our tACS setup was simulated by a simNIBs pipeline based on the 2 mA peak-to-peak intensity of the current on a sample brain. All electrodes were modeled as a 2mm thick rubber layer with a conductivity of 0.1 S/m and the conductive gel was set as 1mm thick with a conductivity of 3 S/m. We normalized the electric field distribution mesh and transferred it to nifty. The modelled current flow in individuals suggests that the extent of the electric field yielded by this 4 × 1 electrode montage is confined to the region of the preSMA (Figure 4 C).

### tACS phase-locked stop signal presentation

In order to understand how phase modulates inhibition, we aligned the stop signal onset with the 4 (Exp 1) and 8 (Exp 2) desired phases of the entraining tACS oscillation as shown in Figure 4 A. The experimental setup of the phase-locked stimulus presentation system was previously accomplished and described in ten Oever et al. (2016). In the setup, source files created in MATLAB (TheMathWorks Inc., Natick, Massachusetts, USA) containing values oscillating between -1 and +1 are loaded into DataStreamer, a custom PC software (https://osf.io/h6b8v/). DataStreamer scales the predetermined tACS waveform signals to the desired tACS intensity and feeds them through a digital-analog converter (DAC) from National Instruments (Austin, TX, USA). A standard BNC cable connected the DAC to the remote input connection of the tACS stimulator. The resolution of the tACS waveform in the source files was 10000 Hz. In a secondary timeline in the source file, stimulus triggering pulses were coded to indicate the timing (by their position in the timeline with respect to the tACS waveform) and parameters (by their numerical value). The digital values were communicated by DataStreamer via the DAC to the PC running the stop signal task in Labview which continuously detected incoming triggering pulses. The pulses triggered the start of each trial and the presentation of the stop signals (stopping of indicator before reaching the target line). The pulses for the stop signal were aligned to the desired phase on the tACS waveform closest to the desired stop signal delay. We quantified the delay between the trigger sent out and the timestamps of the actual stop signal occurrence provided by the task software. The mean of the absolute delay in the 95% range (2.5th to 97.5th percentile) was 1.11 ms and within 2.86 ms, which was calculated per participant and per phase condition (Table S3 and Table S4). Note that these values are additionally constrained to the frame rate of the LCD monitor which was 240 Hz (4.16 ms per frame). When the program received a stop trigger, the actual stop signal would be presented at the next frame (the delay is between 0 and 4.16 ms). The presentation frame rate constrained the number of possible phase bins (20 Hz is 50 ms/cycle, for 8 phase bins, 6.25 ms/bin), but was still acceptable for an 8-phase condition.

### EEG procedures

In experiment 2, EEG signals were recorded through actiCHamp Plus amplifier (Brain Products, Germany) from 8 Ag/AgCl electrodes (Fz, FC1, FC2, C3, Cz; A1, A2 as the references; and AFz as the ground). The sampling rate was 1000 Hz and the impedance was kept below 20 kΩ. In addition, an electrooculogram (EOG) was recorded from the electrodes located at the corner of both eyes as well as both above and below the left eye.

### Behavioral analysis

MATLAB 2022a (Mathworks, Natick, MA, USA) was used to analyze behavioral data. The force data collected trial-by-trial were converted from Volt to Newton. Next, a fifth-order 20 Hz low pass Butterworth filter was applied to filter the force data. In each trial, the average force of the data between -650 ms and -400 ms before the target time was subtracted to correct the baseline. After preprocessing the force data, we calculated the following for all stop trials: (1) peak force, the maximum force of the trial; (2) peak force rate, which was the maximum rate of change in the force signal; The force measures above that were larger than 2.5 × SD of their mean value were seen as the outliers values and deleted. For both the peak force and the peak force rate, we determine the median per phase condition as the distributions were strongly left skewed.

Reaction time (RT) was computed based on the time that the response threshold was initially reached (typically 25% of MVF). We determined the mean RT on failed stops (RTsf) and RT on go trials (RTgo). The go trials with an RT<400 ms or RT>990 ms, identified as early response or no response errors, were removed from the dataset. To be able to calculate the Stop Signal Reaction Time (SSRT), the dataset had to satisfy the assumption of the independent race model (Verbruggen et al., 2019) (i.e., go trials and stop trials needed to behave independently). We compared the mean RTgo against the mean RTsf to see if the former was larger than the latter. The SSRT was then calculated using integration methods (Verbruggen et al., 2019).

The percent of successful stops (P(inhibition)) for each phase condition was calculated based on all five stop signal delays (SSD).

### EEG analysis

The resting EEG data were analyzed using FieldTrip (Oostenveld et al., 2011), in MATLAB 2022a. We pre-processed data by: (1) re-referencing data of each channel to the average of left and right reference channels (A1, A2); (2) removing linear trend and baseline correcting; (3) filtering with a 1-300 Hz band pass filter and a 50 Hz FIR band stop filter; (4) Regressing out eye movements using the function *scrls_regression* of the Eeglab plugin AAR (Gomez-Herrero, 2007) (filter order 3; forgetting factor 0.999; sigma 0.01; precision 50). Data from both pre and post stimulation periods were segmented into 1 s epochs. The epochs including noisy signals were rejected by visual detection. Then, the EEG data underwent a fast Fourier transformation (FFT) in a 4-45 Hz frequency range and a Hanning window with a 10s zeros-padding. Subsequently, the mean absolute beta power at 13-30 Hz was determined and compared between pre and post stimulation.

For the purpose of investigating the correlation between subjects’ individual beta frequency (IBF) and the degree of phase modulation on performance, the averaged pre-resting EEG spectra were log-transformed and fitted by a linear trend in a least-squares manner to remove the 1/f property of the spectra (Haegens et al., 2014; Nikulin and Brismar, 2006). The IBF was then obtained by finding the peak from a 3^*rd*^ order Gaussian curve fitted to the EEG spectra.

### Statistical analysis

All analyses were carried out using MATLAB 2022a and IBM SPSS Statistics 27. In order to validate the hypothesis that inhibition performance is influenced by phase conditions and oscillates along with neuronal oscillations entrained by tACS, we investigated: (a) the presence of tACS-phase effects, and (b) a one-cycle sinus matched the pattern of behavioral outcomes over the sampled phase conditions. Commencing the analysis, a repeated measures ANOVA was performed to assess the difference of stimulation-related discomfort (VAS*discomfort*) and fatigue (VAS*fatigue*) between sessions (Exp 2) and time (pre- and post-stimulation) on the subjective level.

For both Exp 1 and Exp 2, we evaluated the impact of the beta tACS phase on inhibition performance by comparing the outcome measures (SSRT, percent of successful inhibition, peak force and peak force rate) on different phase conditions (4 phase conditions in Exp 1 and 8 in Exp 2). The individual outcome measures were averaged per phase. Repeated measures ANOVA was used to analyze SSRT, RTsf, and P(inhibition), while the Friedman test was employed for assessing the significance of differences in non-normally distributed peak force and peak force rate (Shapiro-Wilk test p<0.05). Post-hoc tests were conducted to identify specific phases that significantly influenced inhibition behavior in terms of SSRT, RTsf, and P(inhibition). Wilcoxon Signed Rank test was used to assess peak force and peak force rate. On both the group level and the individual level, a performance pattern yielded by the outcome measures over tACS oscillatory phase conditions was fitted with a single sinusoidal cycle by the Matlab function ‘fminsearch’ where parameters of the best-fitting phase and amplitude were determined based on minimization of squared errors (Fiebelkorn et al., 2011; ten Oever and Sack, 2015a; Schilberg et al., 2018; de Graaf et al., 2020). The explained variance (R^2^) of the fitted pattern was obtained to evaluate the goodness of fit. The calculation of the R^2^ was based on values at the 4 or 8 sample points. Inspired by previous reports we multiplied R^2^ by the variance of the best fitting sinusoid to obtain ‘relevance values’ (Fiebelkorn et al., 2011; ten Oever and Sack, 2015b; Schilberg et al., 2018; de Graaf et al., 2020). As we had so few sample points, we calculated the variance of the best fitting sinusoid not only based on the sinusoid values at the sample points but calculated population variance of 100 equidistant values on one full cycle of the best fitting sinusoid. This variance measure of the full sinusoid was multiplied with R^2^, which was calculated as 1–(SSM/SST), where SST was the sum of squared differences between observed data at sample points and their mean, and SSM was the sum of squared differences between observed data at sample points and the sinusoid values at those sample points. The resulting hybrid measure of relevance value should reflect both the goodness of fit (R^2^) and the extent of modulation of performance by tACS (de Graaf et al., 2020). The relevance value was then used to perform a permutation test and quantify the related p-values. Conducting the 2000-iteration permutation test, we built a null distribution with 2,000 relevance values that came from the same processing procedure as above, but with the label (phase)-shuffled data. In this way, the null distribution represented the case where the beta phase had no quantitative effect on relevance values. We compared the obtained relevance value against the null distribution and the reported p-values reflect the proportion of the label-shuffled values that have a test statistic larger than that of the non-shuffled data. The relevance value would be deemed statistically significant if it is located within the last 0.05 of the null distribution.

The method described above assumes that individual results are phase-aligned. For various reasons (e.g., transmission time, cortical anatomy), it is not a priori expected that, even in the case of successful tACS entrainment, individual results should phase-align. Considering that mismatched optimal phases might cancel the values of each other out on the group level, we also performed an analysis in which we aligned phases per individual before averaging. Since the phases of individuals were calibrated by shifting the maximum value (peak) of the sinusoidal pattern to the same position (slot 1 in Exp 1 and slot 2 in Exp 2), the statistical results on slot 1 in Exp 1 and slot 2 in Exp 2 should not be considered for subsequent discussions.

## Supplementary information

**Table S1.** Experiment 1 VAS and discomfort rating score.

|  | Pre/Post | Subjective rating between 1-10 | Main effect of time (pre, post) |
| --- | --- | --- | --- |
| Fatigue | Pre | 2.44±1.80 | F(1,28)=1.073<br>p=0.003 |
|  | Post | 3.80±1.95 |  |
| Discomfort | Post | 3.46±0.28 | - |

**Table S2.** Experiment 2 VAS and discomfort rating score.

| Subjective rating between 1-10 |  | Session 1 | Session 2 | Statistic |  |
| --- | --- | --- | --- | --- | --- |
|  |  |  |  | Main effect of phase sessions | Main effect of time (pre, post) |
| Discomfort |  | 3.01±0.19 | 2.57±1.36 | F(1,14)=0.41 p=0.064 | - |
| Fatigue | Pre | 2.62±2.15 | 2.83±2.04 | F(1,14)=0.02 p=0.700 | F=2.54p=0.000 |
|  | Post | 4.15±2.00 | 4.54±2.36 | F(1,14)=0.04 p=0.538 |  |

**Table S3.**
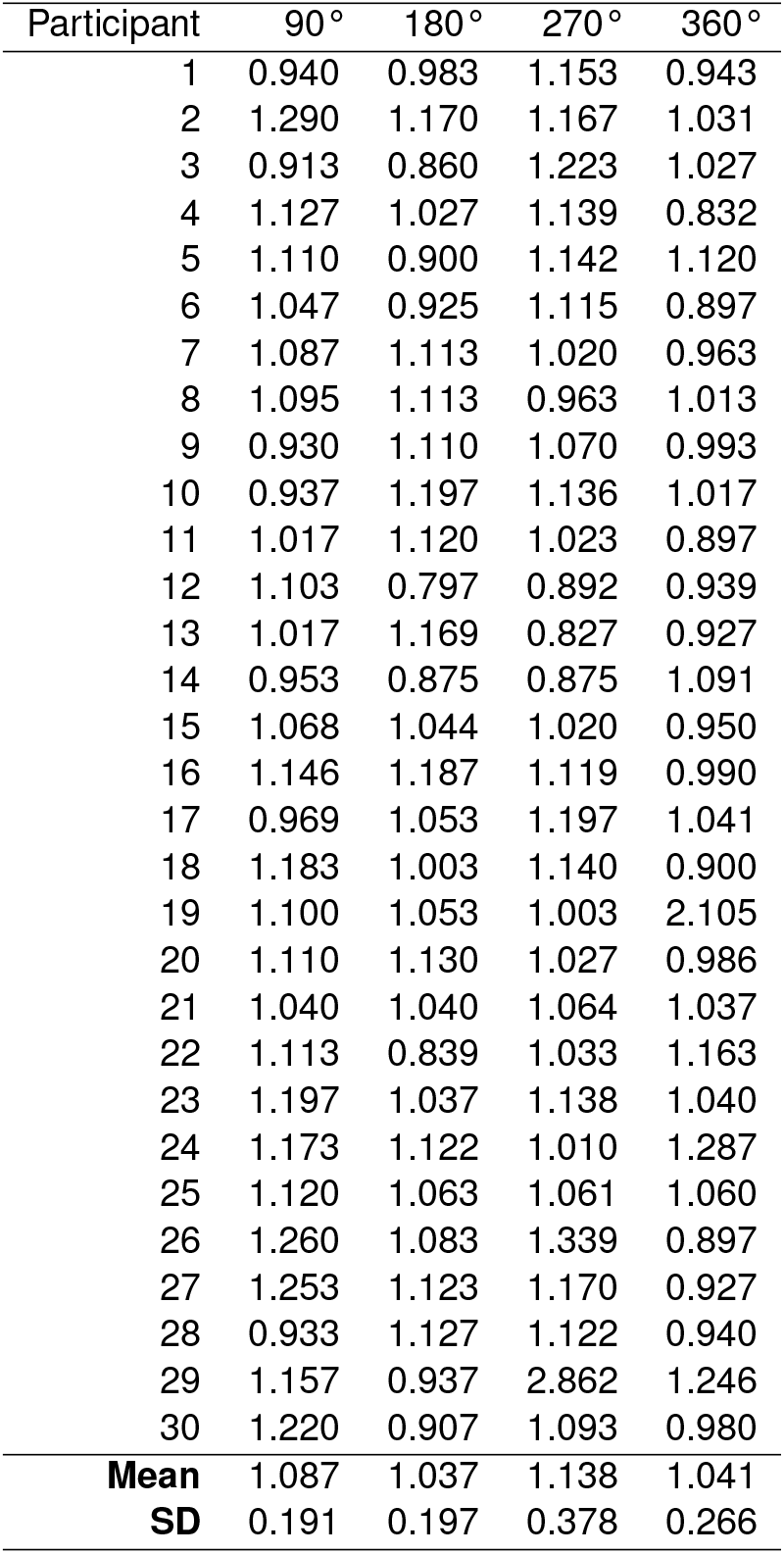
Experiment 1 95% range values of recorded stimulus trigger to stimulus delays, in milliseconds, per phase and participant.

| Participant | 90° | 180° | 270° | 360° |
| --- | --- | --- | --- | --- |
| 1 | 0.940 | 0.983 | 1.153 | 0.943 |
| 2 | 1.290 | 1.170 | 1.167 | 1.031 |
| 3 | 0.913 | 0.860 | 1.223 | 1.027 |
| 4 | 1.127 | 1.027 | 1.139 | 0.832 |
| 5 | 1.110 | 0.900 | 1.142 | 1.120 |
| 6 | 1.047 | 0.925 | 1.115 | 0.897 |
| 7 | 1.087 | 1.113 | 1.020 | 0.963 |
| 8 | 1.095 | 1.113 | 0.963 | 1.013 |
| 9 | 0.930 | 1.110 | 1.070 | 0.993 |
| 10 | 0.937 | 1.197 | 1.136 | 1.017 |
| 11 | 1.017 | 1.120 | 1.023 | 0.897 |
| 12 | 1.103 | 0.797 | 0.892 | 0.939 |
| 13 | 1.017 | 1.169 | 0.827 | 0.927 |
| 14 | 0.953 | 0.875 | 0.875 | 1.091 |
| 15 | 1.068 | 1.044 | 1.020 | 0.950 |
| 16 | 1.146 | 1.187 | 1.119 | 0.990 |
| 17 | 0.969 | 1.053 | 1.197 | 1.041 |
| 18 | 1.183 | 1.003 | 1.140 | 0.900 |
| 19 | 1.100 | 1.053 | 1.003 | 2.105 |
| 20 | 1.110 | 1.130 | 1.027 | 0.986 |
| 21 | 1.040 | 1.040 | 1.064 | 1.037 |
| 22 | 1.113 | 0.839 | 1.033 | 1.163 |
| 23 | 1.197 | 1.037 | 1.138 | 1.040 |
| 24 | 1.173 | 1.122 | 1.010 | 1.287 |
| 25 | 1.120 | 1.063 | 1.061 | 1.060 |
| 26 | 1.260 | 1.083 | 1.339 | 0.897 |
| 27 | 1.253 | 1.123 | 1.170 | 0.927 |
| 28 | 0.933 | 1.127 | 1.122 | 0.940 |
| 29 | 1.157 | 0.937 | 2.862 | 1.246 |
| 30 | 1.220 | 0.907 | 1.093 | 0.980 |
| **Mean** | 1.087 | 1.037 | 1.138 | 1.041 |
| **SD** | 0.191 | 0.197 | 0.378 | 0.266 |

**Table S4.** Experiment 2 95% range values of recorded stimulus trigger to stimulus delays, in milliseconds, per phase and participant.

| Participants | 45° | 90° | 135° | 180° | 225° | 270° | 315° | 360° |
| --- | --- | --- | --- | --- | --- | --- | --- | --- |
| 1 | 1.000 | 1.146 | 1.007 | 1.187 | 1.223 | 1.119 | 0.766 | 0.990 |
| 2 | 1.127 | 1.183 | 1.075 | 1.003 | 1.053 | 1.140 | 1.373 | 0.900 |
| 3 | 1.063 | 1.100 | 1.166 | 1.053 | 0.973 | 1.003 | 1.227 | 2.105 |
| 4 | 1.107 | 1.110 | 1.197 | 1.130 | 1.034 | 1.027 | 1.070 | 0.986 |
| 5 | 1.197 | 1.040 | 0.900 | 1.040 | 1.010 | 1.064 | 1.133 | 1.037 |
| 6 | 2.396 | 1.113 | 2.350 | 0.839 | 2.098 | 1.033 | 2.119 | 1.163 |
| 7 | 1.147 | 1.197 | 1.040 | 1.037 | 1.149 | 1.138 | 1.027 | 1.040 |
| 8 | 1.186 | 1.173 | 1.010 | 1.122 | 1.115 | 1.010 | 0.875 | 1.287 |
| 9 | 1.054 | 1.120 | 1.112 | 1.063 | 0.953 | 1.061 | 1.167 | 1.060 |
| 10 | 0.997 | 1.260 | 0.897 | 1.083 | 1.013 | 1.339 | 1.007 | 0.897 |
| 11 | 0.820 | 1.253 | 1.107 | 1.123 | 0.993 | 1.170 | 0.987 | 0.927 |
| 12 | 1.159 | 0.933 | 1.117 | 1.127 | 1.077 | 1.122 | 1.153 | 0.940 |
| 13 | 1.085 | 1.157 | 1.132 | 0.937 | 1.247 | 2.862 | 1.092 | 1.246 |
| 14 | 1.159 | 1.220 | 1.037 | 0.907 | 1.210 | 1.093 | 1.037 | 0.980 |
| **Mean** | 1.178 | 1.143 | 1.153 | 1.047 | 1.154 | 1.227 | 1.145 | 1.111 |
| **SD** | 0.339 | 0.080 | 0.331 | 0.090 | 0.268 | 0.445 | 0.295 | 0.289 |

**Table S5.** Post hoc tests results of P(inhibition), SSRT and RTsf in Exp. 1.

|  |  | P(inhibition) |  |  | SSRT |  |  | RTsf |  |  |
| --- | --- | --- | --- | --- | --- | --- | --- | --- | --- | --- |
|  |  | Mean diff. | SE | Sig. | Mean diff. | SE | Sig. | Mean diff. | SE | Sig. |
| 90° | 180° | 0.517 | 0.864 | 1.000 | -4.638 | 4.999 | 1.000 | 3.025 | 1.479 | 0.303 |
|  | 270° | -5.632** | 0.966 | 0.000 | 20.000** | 5.402 | 0.006 | 8.491** | 1.527 | 0.000 |
|  | 360° | -7.471** | 1.212 | 0.000 | 17.914* | 6.160 | 0.042 | -0.720 | 1.686 | 1.000 |
| 180° | 270° | -6.149** | 0.880 | 0.000 | 24.638** | 3.898 | 0.000 | 5.466* | 1.734 | 0.023 |
|  | 360° | -7.989** | 1.038 | 0.000 | 22.552** | 3.417 | 0.000 | -3.744 | 2.107 | 0.519 |
| 270° | 360° | -1.839 | 1.070 | 0.580 | -2.086 | 4.311 | 1.000 | -9.210** | 1.930 | 0.000 |

**Table S6.** Wilcoxon signed-rank test results of peak force and peak force rate of stop trials in Exp. 1.

|  |  | Peak force |  | Peak force rate |  |
| --- | --- | --- | --- | --- | --- |
|  |  | Z | Sig. | Z | Sig. |
| 90° | 180° | 1.027 | 0.304 | 1.027 | 0.304 |
|  | 270° | -2.995** | 0.003 | -2.152* | 0.031 |
|  | 360° | -3.189** | 0.001 | -2.843** | 0.004 |
| 180° | 270° | -4.206** | 0.000 | -3.860** | 0.000 |
|  | 360° | -4.271** | 0.000 | -4.054** | 0.000 |
| 270° | 360° | -2.281* | 0.023 | -2.281* | 0.023 |

**Table S7.** Post hoc tests results of P(inhibition), SSRT and RTsf in Exp. 2.

|  |  | P(inhibition) |  |  | SSRT |  |  | RTsf |  |  |
| --- | --- | --- | --- | --- | --- | --- | --- | --- | --- | --- |
|  |  | Mean Diff. | SE | Sig. | Mean Diff. | SE | Sig. | Mean Diff. | SE | Sig. |
| 45 | 90 | 2.565 | 1.723 | 1.000 | -7.821 | 7.597 | 1.000 | -2.458 | 3.246 | 1.000 |
|  | 135 | 2.149 | 2.234 | 1.000 | -22.429 | 10.014 | 1.000 | -4.404 | 2.979 | 1.000 |
|  | 180 | 2.922 | 2.020 | 1.000 | -9.429 | 6.059 | 1.000 | 1.548 | 3.137 | 1.000 |
|  | 225 | 3.160 | 2.920 | 1.000 | -13.679 | 13.099 | 1.000 | 2.396 | 4.063 | 1.000 |
|  | 270 | -2.197 | 2.230 | 1.000 | 14.464 | 7.468 | 1.000 | 7.940 | 3.026 | 0.589 |
|  | 315 | 0.184 | 1.896 | 1.000 | 3.857 | 10.522 | 1.000 | -0.290 | 2.532 | 1.000 |
|  | 360 | -5.888 | 2.319 | 0.692 | 11.107 | 6.620 | 1.000 | -1.653 | 3.367 | 1.000 |
| 90 | 135 | -0.416 | 2.145 | 1.000 | -14.607 | 11.315 | 1.000 | -1.946 | 2.333 | 1.000 |
|  | 180 | 0.357 | 1.080 | 1.000 | -1.607 | 7.473 | 1.000 | 4.006 | 2.117 | 1.000 |
|  | 225 | 0.595 | 2.647 | 1.000 | -5.857 | 12.495 | 1.000 | 4.854 | 3.376 | 1.000 |
|  | 270 | -4.762 | 1.317 | 0.088 | 22.286 | 8.684 | 0.657 | 10.398** | 2.164 | 0.010 |
|  | 315 | -2.381 | 1.762 | 1.000 | 11.679 | 9.714 | 1.000 | 2.168 | 3.270 | 1.000 |
|  | 360 | -8.452* | 1.941 | 0.022 | 18.929 | 10.003 | 1.000 | 0.805 | 2.224 | 1.000 |
| 135 | 180 | 0.773 | 2.169 | 1.000 | 13.000 | 9.157 | 1.000 | 5.952 | 2.691 | 1.000 |
|  | 225 | 1.011 | 2.550 | 1.000 | 8.750 | 10.096 | 1.000 | 6.801 | 2.292 | 0.305 |
|  | 270 | -4.346 | 2.119 | 1.000 | 36.893* | 8.665 | 0.026 | 12.344* | 2.828 | 0.021 |
|  | 315 | -1.965 | 2.135 | 1.000 | 26.286 | 10.043 | 0.596 | 4.114 | 2.257 | 1.000 |
|  | 360 | -8.037 | 2.558 | 0.218 | 33.536 | 10.015 | 0.147 | 2.751 | 3.384 | 1.000 |
| 180 | 225 | 0.238 | 3.100 | 1.000 | -4.250 | 11.970 | 1.000 | 0.849 | 3.793 | 1.000 |
|  | 270 | -5.119* | 1.139 | 0.017 | 23.893 | 6.524 | 0.080 | 6.392 | 2.446 | 0.600 |
|  | 315 | -2.738 | 2.387 | 1.000 | 13.286 | 9.392 | 1.000 | -1.838 | 3.328 | 1.000 |
|  | 360 | -8.810** | 1.435 | 0.001 | 20.536 | 5.353 | 0.058 | -3.201 | 3.131 | 1.000 |
| 225 | 270 | -5.357 | 2.622 | 1.000 | 28.143 | 9.803 | 0.368 | 5.544 | 3.972 | 1.000 |
|  | 315 | -2.976 | 1.817 | 1.000 | 17.536 | 13.555 | 1.000 | -2.686 | 2.540 | 1.000 |
|  | 360 | -9.048 | 3.291 | 0.463 | 24.786 | 13.537 | 1.000 | -4.049 | 4.420 | 1.000 |
| 270 | 315 | 2.381 | 2.159 | 1.000 | -10.607 | 9.435 | 1.000 | -8.230 | 3.419 | 0.887 |
|  | 360 | -3.690 | 1.367 | 0.509 | -3.357 | 6.747 | 1.000 | -9.593 | 2.789 | 0.123 |
| 315 | 360 | -6.071 | 2.548 | 0.927 | 7.250 | 9.424 | 1.000 | -1.363 | 3.623 | 1.000 |

**Figure S1.**
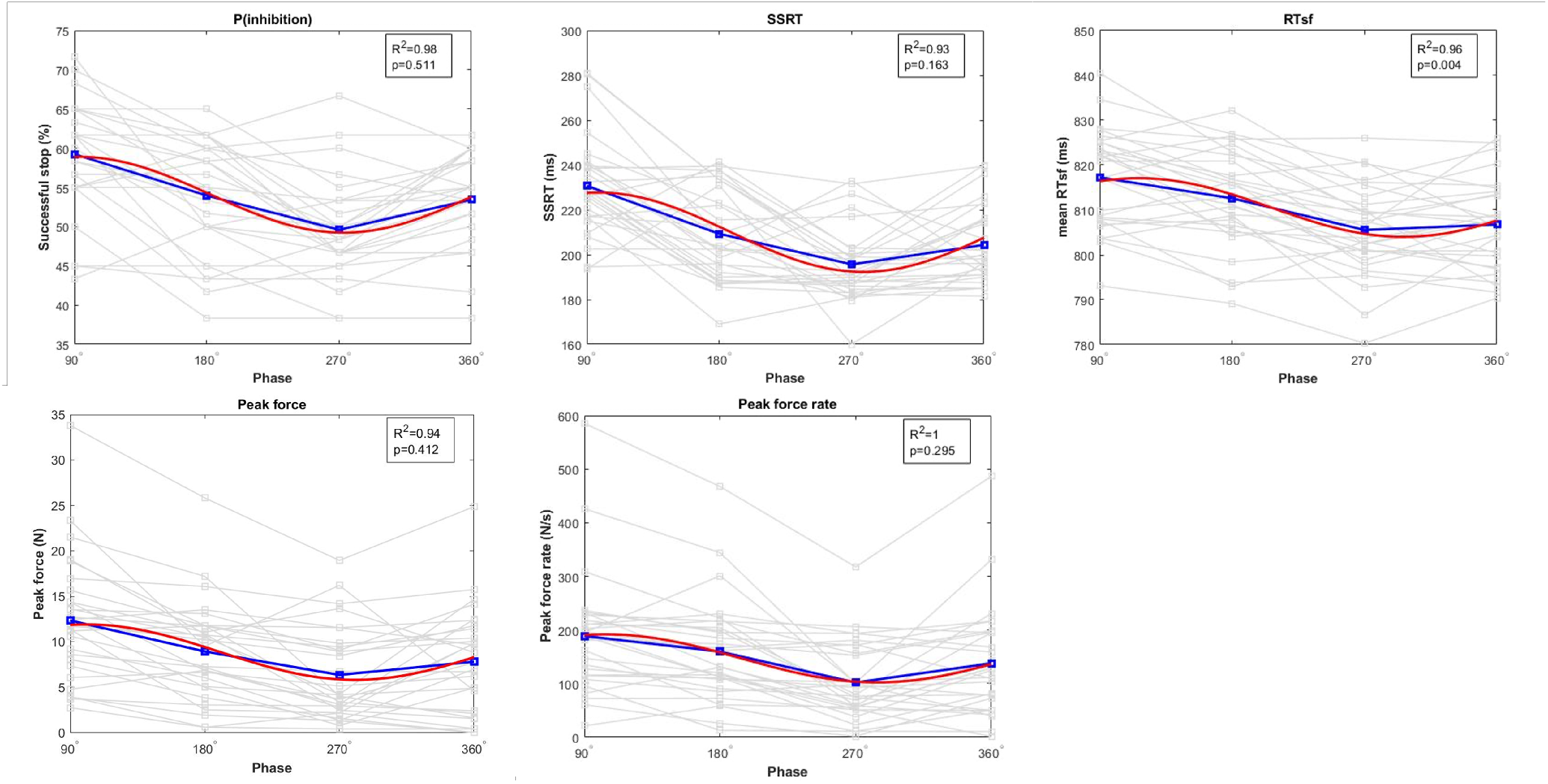
Phase-aligned group results of Exp.1

**Table S8.** Wilcoxon signed-rank test results of peak force and peak force rate of stop trials in Exp. 2.

|  |  | Peak force |  | Peak force rate |  |
| --- | --- | --- | --- | --- | --- |
|  |  | Z | Sig. | Z | Sig. |
| 45 | 90 | 1.528 | 0.124 | 1.601 | 0.109 |
|  | 135 | 1.601 | 0.109 | 1.664 | 0.096 |
|  | 180 | 1.475 | 0.140 | 1.852 | 0.064 |
|  | 225 | 1.601 | 0.109 | 2.229* | 0.026 |
|  | 270 | 0.973 | 0.331 | 1.036 | 0.300 |
|  | 315 | -0.094 | 0.925 | 0.282 | 0.778 |
|  | 360 | -0.973 | 0.331 | -0.659 | 0.510 |
| 90 | 135 | 0.031 | 0.975 | -0.785 | 0.433 |
|  | 180 | 1.161 | 0.245 | 1.664 | 0.096 |
|  | 225 | -0.094 | 0.925 | -0.094 | 0.925 |
|  | 270 | -2.471* | 0.016 | -1.601 | 0.109 |
|  | 315 | -1.977* | 0.048 | -1.412 | 0.158 |
|  | 360 | -2.982** | 0.003 | -2.542* | 0.011 |
| 135 | 180 | -0.345 | 0.730 | 1.224 | 0.221 |
|  | 225 | -0.094 | 0.925 | 0.534 | 0.594 |
|  | 270 | -2.229* | 0.026 | -1.036 | 0.300 |
|  | 315 | -2.040* | 0.041 | -1.789 | 0.074 |
|  | 360 | -2.919** | 0.004 | -2.605* | 0.009 |
| 180 | 225 | -0.220 | 0.826 | -0.534 | 0.594 |
|  | 270 | -3.296** | 0.001 | -2.982** | 0.003 |
|  | 315 | -1.852 | 0.064 | -2.354* | 0.019 |
|  | 360 | -3.170* | 0.002 | -3.107** | 0.002 |
| 225 | 270 | -1.475 | 0.140 | -1.475 | 0.140 |
|  | 315 | -2.480* | 0.013 | -2.480* | 0.013 |
|  | 360 | -2.354* | 0.019 | -2.542* | 0.011 |
| 270 | 315 | -0.220 | 0.826 | -0.659 | 0.510 |
|  | 360 | -2.354* | 0.019 | -2.345* | 0.019 |
| 315 | 360 | -1.224 | 0.221 | -1.224 | 0.221 |

**Figure S2.**
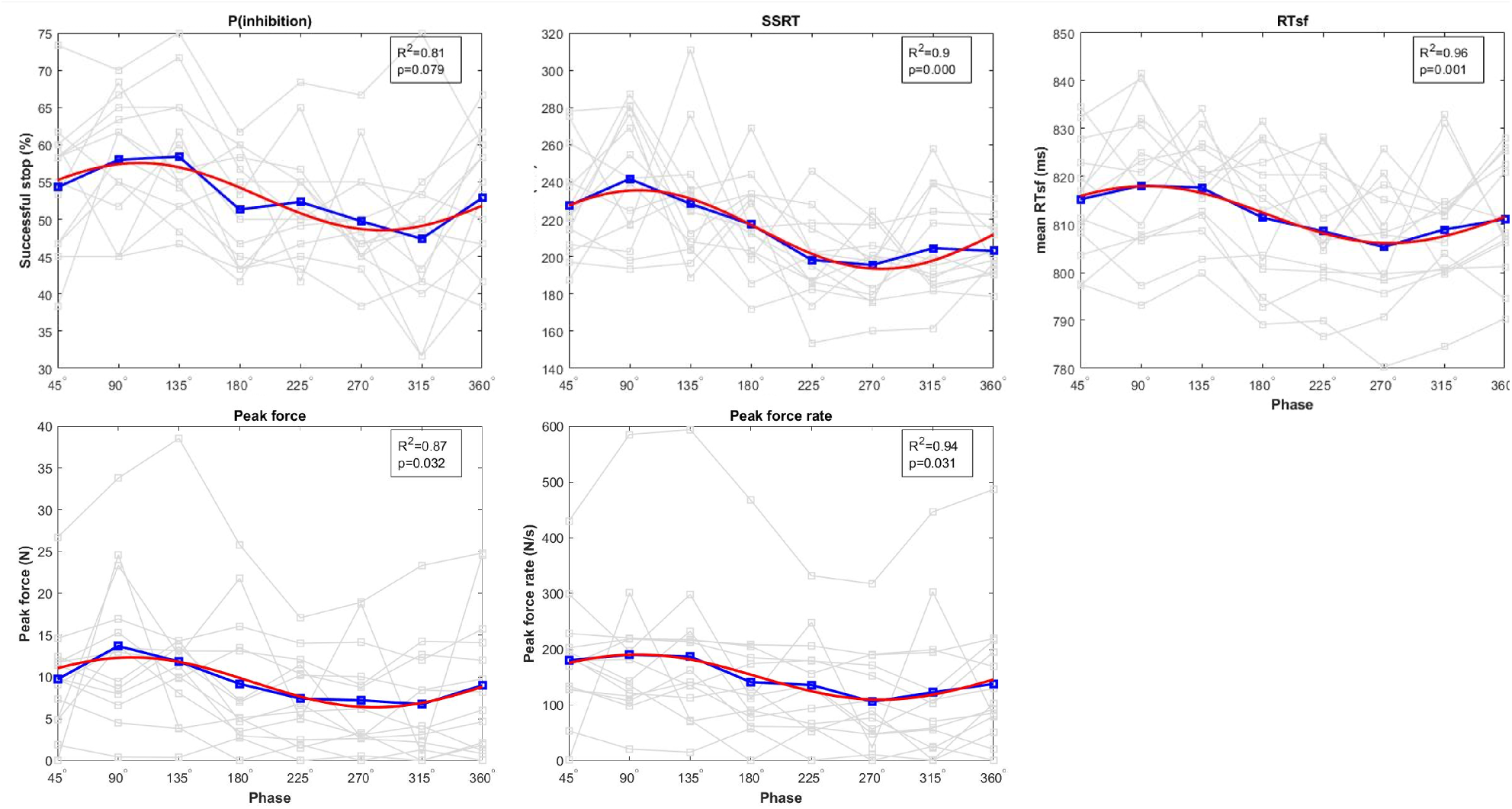
Phase-aligned group results of Exp.2

